# Role of *Pseudomonas aeruginosa* Dnr-regulated denitrification in oxic conditions

**DOI:** 10.1101/2025.03.31.646406

**Authors:** Stacie Stuut Balsam, Dallas L. Mould, Fabrice Jean-Pierre, Deborah A. Hogan

## Abstract

*Pseudomonas aeruginosa* causes acute and chronic infections such as those that occur in the lungs of people with cystic fibrosis (CF). In infection environments, oxygen (O_2_) concentrations are often low. The transcription factor Anr responds to low O_2_ by upregulating genes necessary for *P. aeruginosa* fitness in microoxic and anoxic conditions. Anr regulates *dnr*, a gene encoding a transcriptional regulator that promotes the expression of genes required for using nitrate as an alternative electron acceptor during denitrification. In CF sputum, transcripts involved in denitrification are highly expressed. While Dnr is necessary for the anoxic growth of *P. aeruginosa* in CF sputum and artificial sputum media (ASMi), the contribution of denitrification to *P. aeruginosa* fitness in oxic conditions has not been well described. Here we show that *P. aeruginosa* requires *dnr* for fitness in ASMi and the requirement for *dnr* is abolished when nitrate is excluded from the media. Additionally, we show that *P. aeruginosa* consumes nitrate in lysogeny broth (LB) under microoxic conditions. Furthermore, strains without a functioning quorum sensing regulator LasR, which leads to elevated Anr activity, consume nitrate in LB even in normoxia. There was no growth advantage for *P. aeruginosa* when nitrate was present at concentrations from 100 µM to 1600 µM. However, *P. aeruginosa* consumption of nitrate in oxic conditions created a requirement for Dnr and Dnr-regulated NorCB likely due to the need to detoxify nitric oxide. These studies suggest that Anr- and Dnr-regulated processes may impact *P. aeruginosa* physiology in many common culture conditions.

**Importance:** *Pseudomonas aeruginosa* is an opportunistic pathogen commonly isolated from low-oxygen environments such as the lungs of people with cystic fibrosis. While the importance of *P. aeruginosa* energy generation by denitrification is clear in anoxic environments, the effects of denitrification in oxic cultures is not clear. Here, we show that nitrate is consumed even in oxic environments and while it does not appear to stimulate growth, it does impact fitness. Further, we report that two regulators that are best known for their roles in anoxic conditions also contribute to *P. aeruginosa* fitness in commonly-used laboratory media in presence of oxygen.

## Introduction

*Pseudomonas aeruginosa* is an important opportunistic pathogen often isolated from microoxic environments. For example, within a mucus plug in a lung of an individual with cystic fibrosis (CF), oxygen (O_2_) concentrations are as low as 7 µM (1). Additionally, *P. aeruginosa* often forms biofilms which have steep oxygen gradients with microoxic zones (2, 3). The O_2_-sensitive transcription factor, Anr (**a**naerobic regulation of arginine deiminase and **n**itrate **r**eduction), plays an important role in *P. aeruginosa* adaptation in low O_2_ environments (4–6). Anr activation is dependent on the formation and insertion of an O_2_-labile [4Fe-2S]^2+^ cofactor that is required for Anr dimerization (4, 7). Once active, dimeric Anr induces the expression of many genes relevant to life in anoxic and microoxic conditions (7, 8). When O_2_ is limited, Anr upregulates one of two high-affinity cytochrome c oxidases (*cbb*_3_-1 and *cbb*_3_-2) that allow for aerobic respiration even when concentrations of O_2_ are as low as 3 µM (9). Anr also regulates *mhr* which is epistatic to the *cbb*_3_ oxidases and encodes for a hemerythrin protein that binds O_2_ with micromolar affinities (10, 11).

Anr has been shown to be important in several settings in which O_2_ is present. It is necessary for full fitness of *P. aeruginosa* in colony biofilms (10) and contributes to *P. aeruginosa* biofilm growth in lung surfactant medium (12) and an artificial sputum medium for imaging (ASMi) made to mimic the CF lung environment (13). The *P. aeruginosa* Δ*anr* mutant also has a severe defect in a murine pneumonia model (12). Loss-of-function mutations in the gene encoding the transcription factor LasR are frequently found (14–16) and lead to significantly higher Anr activity than comparable strains with functional LasR (10, 17). This high Anr activity contributes to the competitive fitness advantage of a LasR-strain over its LasR+ comparator (10). While Anr is important in biofilms and in infections, its importance in oxic planktonic cultures has not been well described.

Anr directly regulates the gene encoding for Dnr (**d**issimilative **n**itrate respiration **r**egulator) (18, 19). Dnr is a transcriptional regulator of multiple genes that participate in *P. aeruginosa* denitrification, a process in which nitrate (NO_3_^-^) is used as an alternative electron acceptor through nitrite (NO_2_^-^), nitric oxide (NO), and nitrous oxide (N_2_O) intermediates ultimately leading to the formation of nitrogen gas (N_2_) (20, 21). The Anr-Dnr regulatory cascade is complex with overlap between Anr and Dnr binding sites (22). In addition to its direct regulation by Anr, *dnr* is also regulated by NarXL, a two-component system that, itself, is regulated by Anr (18). NarXL also regulates the *nar* genes that encode the nitrate reductase enzymes (8, 23). Anr, Dnr, and NarXL along with downstream regulators control the expression of the *nir*, *nor* and *nos* genes that encode nitrite reductases, nitric oxide reductases and nitrous oxide reductases, respectively (8, 24–26). Dnr also directly regulates the *nar, nir, nor* and *nos* genes (8, 18, 24, 25).

Transcripts associated with denitrification are highly upregulated in CF clinical isolates and strains of *P. aeruginosa* grown ex vivo, under anoxic conditions in CF sputum (27–29) and laboratory media meant to emulate the CF lung environment like ASMi and synthetic CF sputum media (SCFM and its derivatives) (28, 30, 31). Nitrate concentrations in CF sputum and in media that model sputum are ∼300-400 µM (32). At these concentrations, *P. aeruginosa* predominantly uses aerobic respiration to support growth, but genes involved in denitrification are also upregulated (9). While it is known that denitrification can occur in oxic zones of colony biofilms (33, 34), its effects on *P. aeruginosa* fitness has not been well described.

Herein, we show that a *P. aeruginosa* Δ*anr* mutant is defective in growth in lysogeny broth (LB) due to defects in denitrification. We found ∼130 µM nitrate in LB, mostly from the yeast extract medium component, and found that *P. aeruginosa* consumed the nitrate in LB in microoxic (1% O_2_) but not normoxic (21% O_2_) conditions. LasR-strains consumed nitrate in both normoxia and microoxia. In correlation with nitrate consumption, the Δ*dnr* mutant had a growth defect in 1% O_2_ while a Δ*lasR*Δ*dnr* mutant had a growth defect in LB at 21% and 1% O_2_. Growth defects in Δ*dnr* and Δ*lasR*Δ*dnr* mutants were only observed in tryptone broth (TB) and ASMi, an artificial sputum medium, when nitrate was present. Genetic analysis of mutants defective in denitrification found that nitric oxide (NO) detoxification was the most important process in Dnr-dependent contributions to growth. Together, these data suggest that Dnr was important for fitness even when nitrate is available at micromolar concentrations in the presence of O_2_.

## Results

### Dnr-regulated denitrification contributes to Δ*anr* growth defect in the presence of nitrate

After 16 h of growth in LB at 21% and 1% O_2_, we found that a Δ*anr* strain grew ∼20% less than the WT at both oxygen concentrations (P=0.007), and that complementation of *anr* restored growth back to WT levels (Fig. 1A). Decreased expression of Anr-regulated *mhr*, an O_2_ binding protein shown to be important for competitive fitness in colony biofilms (10, 11), did not explain the lower culture yield as the growth of the WT strain and the Δ*mhr* mutant had no significant differences in microoxic growth (Fig. S1).

**Figure 1.**
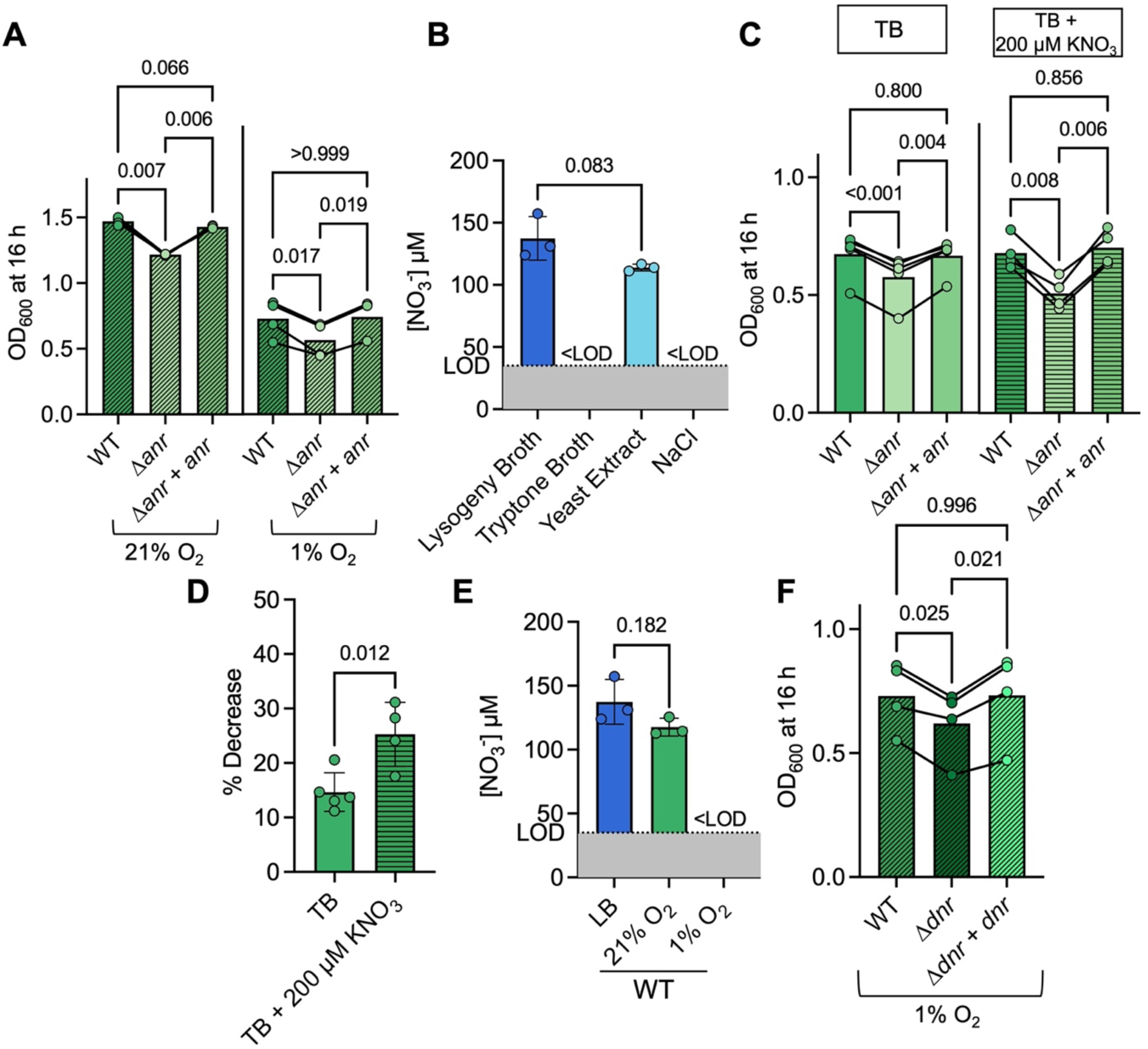
Contribution of Anr and Dnr to *P. aeruginosa* growth in media +/- nitrate. **A.** *P. aeruginosa* strain culture density of PA14 wild type (WT), the Δ*anr* mutant, and the Δ*anr*+*anr* strain after 16 h in LB at 21% and 1% O_2_ in a 96-well plate. **B.** Levels of NO_3_^-^ in lysogeny broth (LB), tryptone broth (TB), a 0.5% yeast extract solution, and a 0.5% sodium chloride (NaCl) solution. The dotted line indicates the lower limit of detection (LOD). **C.** WT, Δ*anr* and Δ*anr*+*anr* culture density after 16 h of growth in TB (solid) or TB + 200 µM KNO_3_ (vertical stripes) at 1% O_2_. Each point represents an average of replicates from one day and lines connect data from the same experiment. **D.** The % decrease in growth of the Δ*anr* strain relative to WT in TB versus TB + 200 µM KNO_3_ in 96-well plates at 1% O_2_ for 16 h with shaking. **E.** The levels of NO_3_^-^ in LB before and after WT growth for 16 h at 21% and 1% O_2_. NO_3_^-^ levels were calculated using a standard curve of KNO_3_ in water and normalized to OD_600_. The levels of nitrate in LB are the same as in panel B. **F.** WT, the Δ*dnr* mutant, and the Δ*dnr+dnr* strain culture density after growth in LB (diagonal stripes) at 1% O_2_ for 16 h in a 96-well plate with shaking. P-values were calculated using a paired one-way ANOVA with multiple comparisons (A, C, and F) and an unpaired t-test (B, D and E).

To determine if differences in denitrification could have contributed to the reduced growth of the Δ*anr* mutant, we first measured nitrate in LB. We found that LB contained ∼130 µM nitrate (Fig. 1B) and negligible levels of nitrite (<5 µM, Fig. S2). LB is composed of tryptone, yeast extract and sodium chloride (NaCl), and analysis of each component showed that the majority of the nitrate was in yeast extract (P=0.083) and levels of nitrate in the tryptone and salt components of LB were both below the limit of detection (Fig. 1B). To study the effects of low concentrations of nitrate on *P. aeruginosa* growth under normoxic and microoxic conditions, we used tryptone broth (TB) as a base medium without or with 200 µM KNO_3_ added. When we grew WT and the Δ*anr* mutant in TB or TB + 200 µM KNO_3_ (Fig. 1C), we found that the percent decrease in growth of the Δ*anr* mutant compared to the WT was significantly greater in TB + 200 µM KNO_3_ (25%±5.9) than in TB (14%±3.7) (Fig. 1D) indicating that differences in denitrification likely contributed to the observed growth defect of the Δ*anr* strain in LB.

To determine if *P. aeruginosa* consumed the nitrate present in LB, we quantified the levels of nitrate in supernatants after 16 h of growth in 5 mL cultures. When grown at 21% O_2_, the levels of nitrate in the PA14 WT supernatant were similar to those in LB, however, at 1% O_2_, the WT strain depleted nitrate to levels below the limit of the detection (Fig. 1E). After growth at 21% O_2_ in 5 mL tube-grown cultures, a condition with no nitrate consumption (Fig. 1E), no significant differences between WT and Δ*dnr* growth were observed (P=0.906). Similarly, only modest differences between WT and Δ*dnr* growth were observed (7% less growth in Δ*dnr*, P=0.037) in 200 µL cultures in 96-well plates incubated with shaking (Fig. S3A). However, in 1% O_2_, a condition in which WT consumed nitrate, the Δ*dnr* mutant had a 15% lower yield (P=0.025) than the WT, and the Δ*dnr*+*dnr* strain restored growth (Fig. 1F).

### *P. aeruginosa* LasR- strains consumes nitrate even in normoxia resulting in Dnr-dependent growth

Like a WT strain, a Δ*lasR* mutant also consumed nitrate in LB to levels below the limit of detection after growth at 1% O_2_ (Fig. 2A). After growth at 1% O_2_ in 200 µL of LB in 96-well plates while shaking, the Δ*lasR*Δ*dnr* mutant had ∼26% less growth when compared to the Δ*lasR* strain (P=0.040) (Fig. 2B). Unlike the WT, which only consumed nitrate in microoxia, the Δ*lasR* mutant also consumed nitrate in LB in cultures grown in 21% O_2_ (Fig. 2A). We observed a similar result with a LasR- strain clinical isolate, J215 (17). In 5 mL LB at 21% O_2_, both J215 and the J215 Δ*dnr* mutant consumed nitrate in LB to levels below the limit of detection (Fig. S4A) and in these conditions, the J215 Δ*dnr* mutant grew ∼15% less than J215 (Fig. S4B).

**Figure 2.**
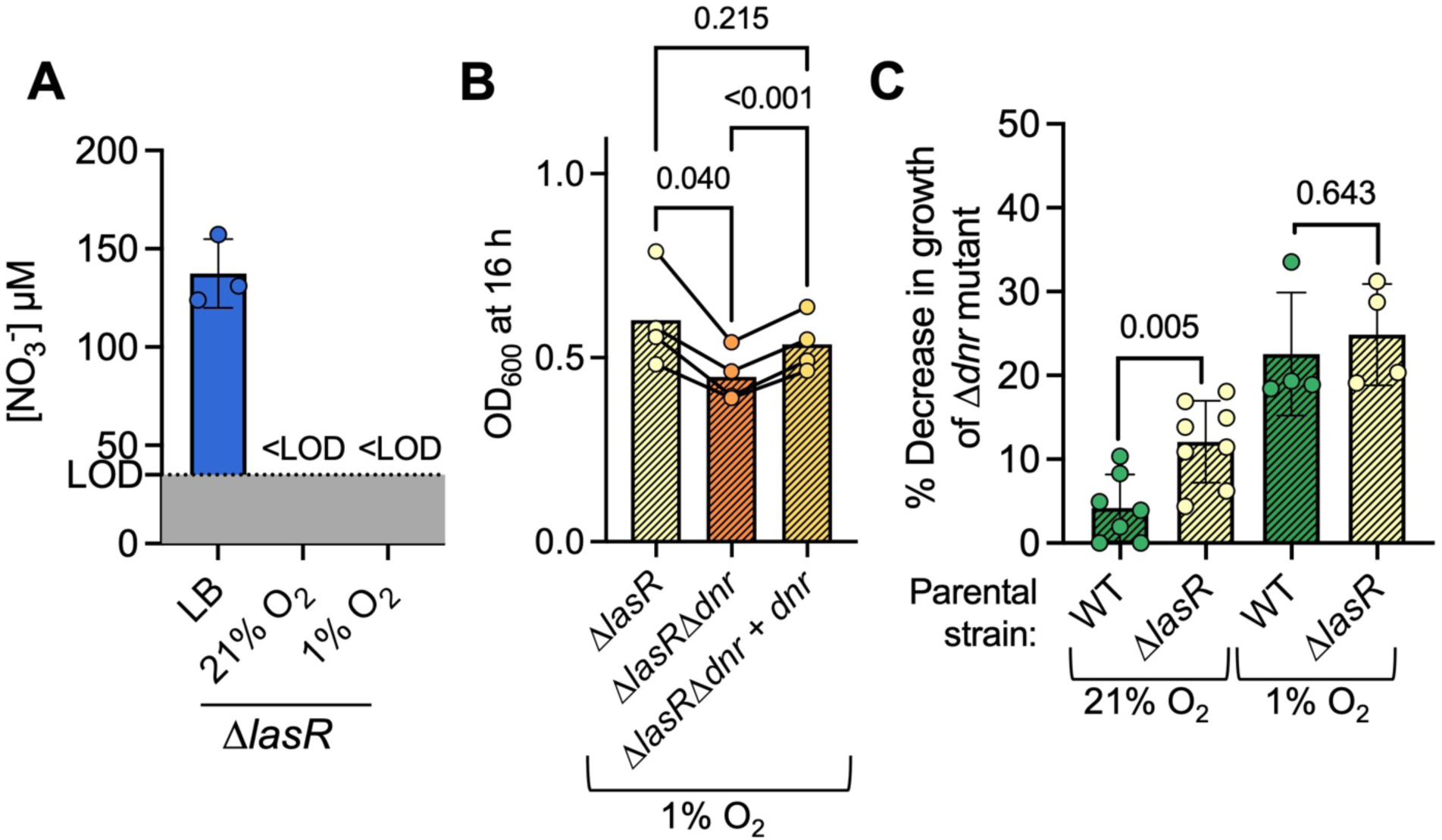
Contribution of Dnr to *P. aeruginosa* growth in concentrations of O_2_ that lead to nitrate consumption. **A.** The levels of NO_3_^-^ in LB before and after Δ*lasR* growth for 16 h at 21% and 1% O_2_. NO_3_^-^ levels were normalized to OD_600_. The levels of nitrate in LB are the same as in Figure 1B. **B.** Δ*lasR*, the Δ*lasR*Δ*dnr* mutant, and the Δ*lasR*Δ*dnr+dnr* strain culture density after growth in LB (diagonal stripes) at 1% O_2_ for 16 h in a 96-well plate with shaking. **C.** The % decrease in growth of the Δ*dnr* and Δ*lasR*Δ*dnr* mutants relative to WT and Δ*lasR* parental strains in LB after 16 h of growth in 96-well plates in 21% or 1% O_2_. Data points represent an average of technical replicates with lines showing comparisons of averages of data from the same day. P-values were calculated using paired one-way ANOVA with multiple comparisons (B) and a paired t-test (C).

After growth in 1% O_2_, when both the WT strain and Δ*lasR* mutant consumed nitrate in LB, the Δ*dnr* and Δ*lasR*Δ*dnr* mutants had a similar % decrease in growth when compared to their parental strains (P=0.643) (Fig. 2C). At 21% O_2_, when only the Δ*lasR* mutant and not the WT consumed nitrate, the percent decrease in growth of a Δ*lasR*Δ*dnr* mutant compared to the Δ*lasR* (∼15%) was significantly higher than a Δ*dnr* mutant compared to the WT (∼4%) (P=0.005) (Fig. 2C). These data show that Δ*dnr* and Δ*lasR*Δ*dnr* growth defects only occurred in cultures in which nitrate was consumed.

### Dnr is important for fitness in TB with nitrate, but does not contribute to increased overall yield of *P. aeruginosa*

Similar to what was observed for the Δ*anr* mutant (Fig. 1C), a Δ*dnr* mutant had a significant growth defect in TB + 200 µM KNO_3_ and not in TB alone (Fig. 3A). In TB with nitrate, WT grew to an average OD of 0.52 ± 0.10 versus Δ*dnr* which grew ∼18% less to an OD of about 0.43 ± 0.10 (Fig. 3A); in contrast WT and Δ*dnr* had similar ODs (0.54 ± 0.07 versus 0.49 ± 0.06, P=0.051) in TB. Similarly, the Δ*lasR*Δ*dnr* mutant grew ∼25% less than the Δ*lasR* parental strain in TB + 200 µM KNO_3_ (P=0.003), while they grew similarly in TB alone (P=0.155) (Fig. 3B). These data further support the model that nitrate consumption was important for the requirement of Dnr for full growth.

**Figure 3.**
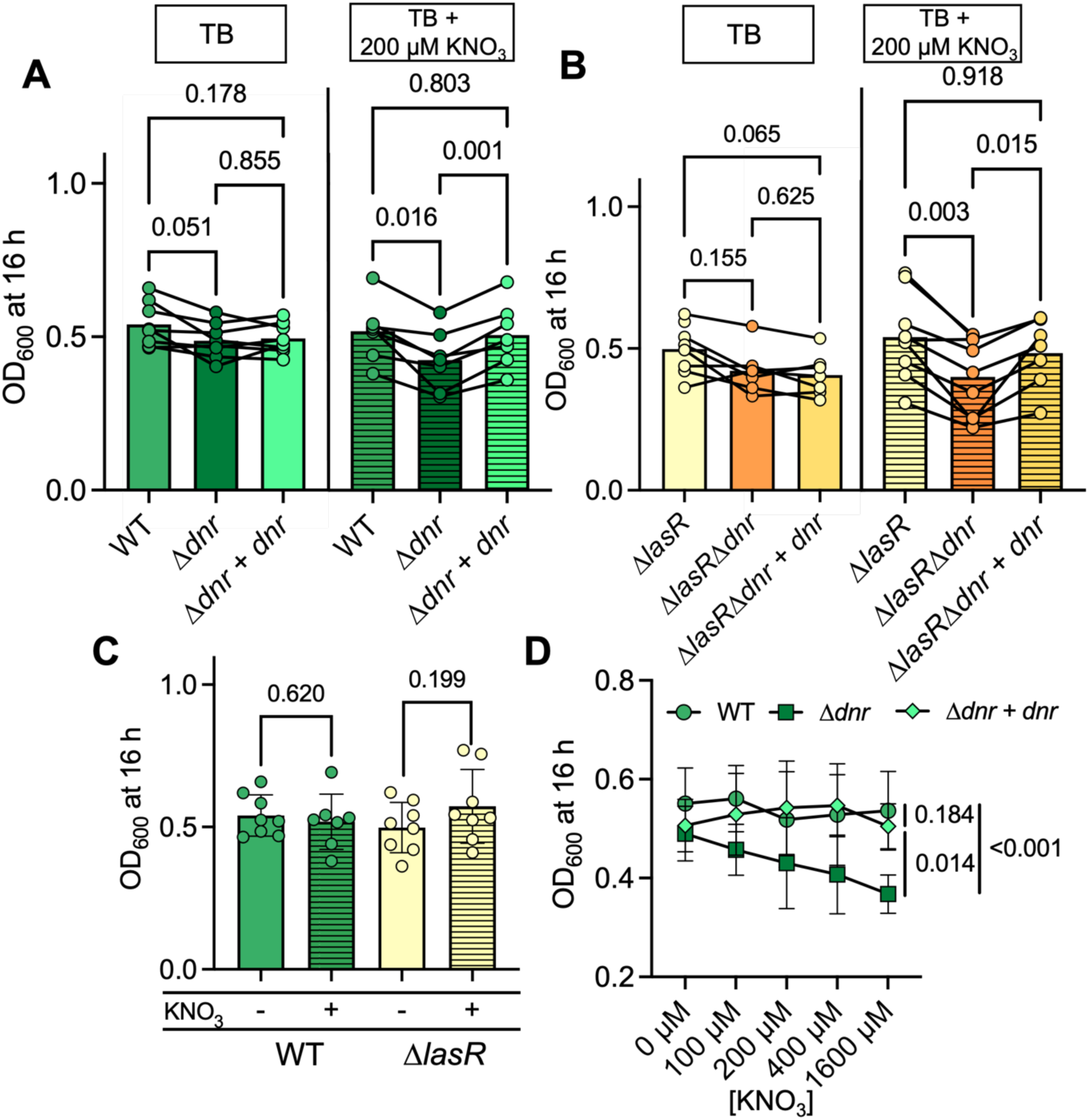
Dnr contribution to microoxic growth and overall yield of *P. aeruginosa* in media +/- nitrate. **A.** WT, the Δ*dnr* mutant, and the Δ*dnr+dnr* strain culture density after growth in TB (solid) and TB + KNO_3_ (horizontal stripes) at 1% O_2_ for 16 h in a 96-well plate with shaking. **B.** Culture densities of Δ*lasR*, a Δ*lasR*Δ*dnr* mutant, and the Δ*lasR*Δ*dnr*+*dnr* strain after 16 h of growth in TB (solid) and TB + 200 µM KNO_3_ (horizontal stripes) in 96-well plates at 1% O_2_. Each data point represents an average of replicates from one day with lines connecting data from the same experiment. **C.** Comparison of growth of WT and Δ*lasR* in TB (solid) and TB + 200 µM KNO_3_ (horizontal stripes) from panels A and B. **D.** Comparison of OD_600_ of WT, Δ*dnr* and Δ*dnr*+*dnr* in TB with 0, 100, 200, 400 and 1600 µM KNO_3_ added. Area under the curve (AUC) was calculated and used for dose response analysis. P-values were calculated using a paired one-way ANOVA (A and B), a paired t-test (C) and a t-test comparison of area under the curve (D).

Interestingly, there was not significantly more growth in TB + 200 µM KNO_3_ than in TB alone for either the WT strain or the Δ*lasR* mutant (Fig. 3C) (WT: 0.54±0.07 in TB and 0.52±0.10 in TB + 200 µM KNO_3_ (P=0.620); Δ*lasR*: 0.50±0.09 in TB + 200 µM KNO_3_ and 0.57±0.13 in TB, P=0.199) (Fig. 3C). Thus, the addition of 200 µM KNO_3_ did not contribute to an increase in the final yield of either strain. Additionally, there was no observed increase in growth for the WT upon the addition of 400 or even 1600 µM KNO_3_. However, as the concentration of KNO_3_ increased, the growth of a Δ*dnr* mutant decreased in a dose-dependent manner (P<0.001) and like WT, there was no dose-response for the Δ*dnr*+*dnr* strain (P=0.184) (Fig. 3D). These data indicate that Dnr was important for growth in the presence of nitrate, but did not lead to an increase in overall yield as more nitrate was made available to consume.

### Dnr was required for fitness in ASMi due to the presence of KNO_3_

We sought to determine if Anr and Dnr contributed to the growth of *P. aeruginosa* WT and Δ*lasR* strains in ASMi, an optically clear version of the synthetic sputum medium SCFM2 (30) which contains 340 µM KNO_3_ (13). In both WT and Δ*lasR* backgrounds, the absence of *anr* caused no significant growth defect in ASMi at 21% O_2_ however, *anr* mutants grew 21% and 15% less than their WT and Δ*lasR* parental strains at 1% O_2_ (Fig. S5A and B). Like for the Δ*anr* mutants, the Δ*dnr* mutants in the WT and Δ*lasR* backgrounds grew significantly less than their parental strains in ASMi in microoxic conditions with ∼25% and ∼30% reduction in growth, respectively, when *dnr* was absent. The microoxic growth differences between WT and Δ*lasR* and their Δ*dnr* derivatives in ASMi was abolished when KNO_3_ was omitted from the medium (Fig. 4A and B). Unexpectedly, Dnr was only necessary for growth in ASMi in 21% O_2_ in the WT background (Fig. S5C), but not in the Δ*lasR* background (Fig. S5D). As in TB (Fig. 3C), the presence of nitrate did not affect culture yield in ASMi for parental strains with functional Dnr (Fig. 4C).

**Figure 4.**
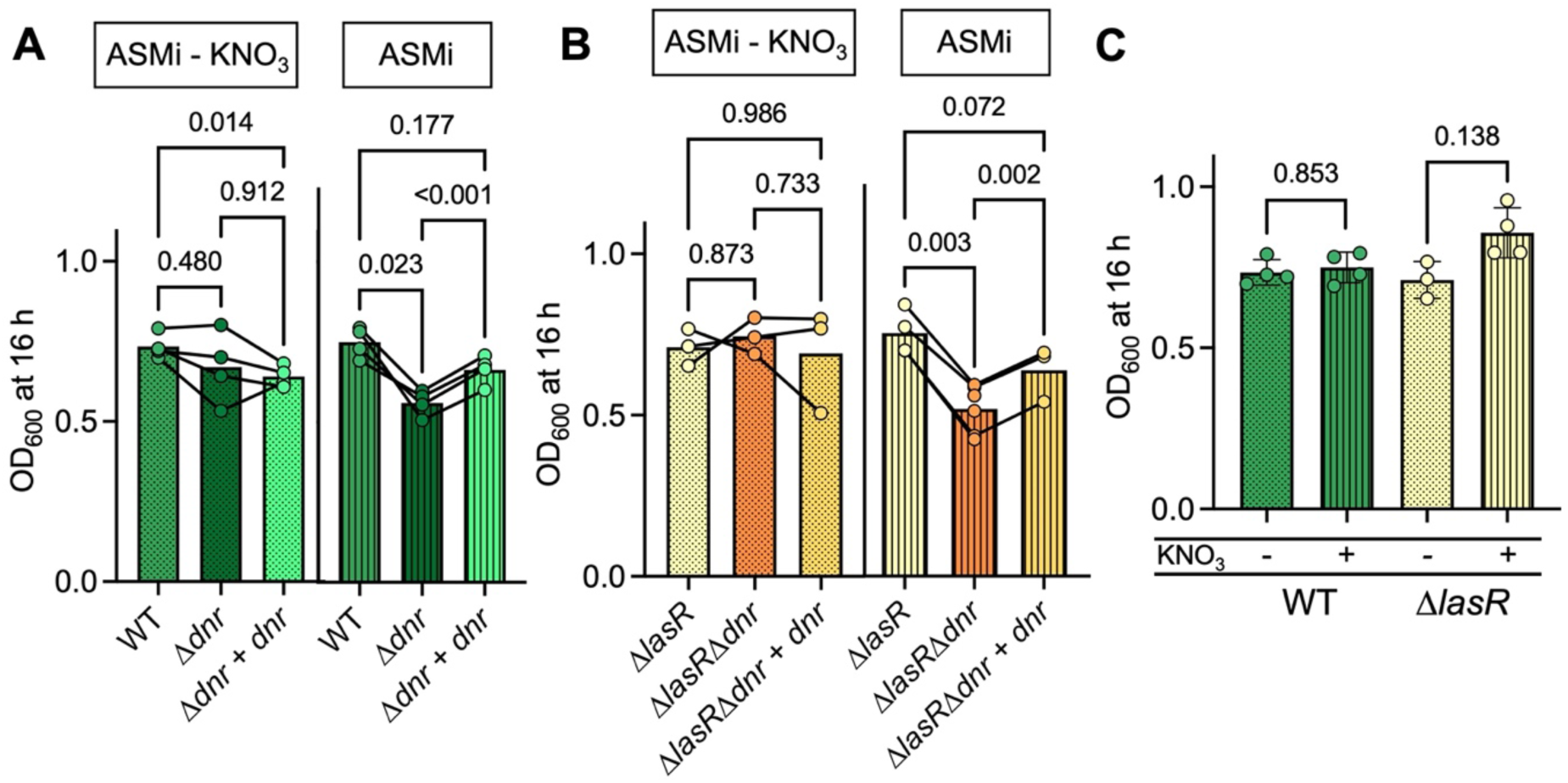
Dnr contribution to microoxic growth of *P. aeruginosa* in ASMi +/- nitrate. **A.** WT, the Δ*dnr* mutant, and the Δ*dnr+dnr* strain culture density after growth in ASMi - KNO_3_ (dots) and ASMi (vertical stripes) at 1% O_2_ for 16 h in a 96-well plate at 1% O_2_ with shaking. **B.** The Δ*lasR*, Δ*lasR*Δ*dnr*, and the Δ*lasR*Δ*dnr+dnr* strain culture densities after growth in ASMi - KNO_3_ (dots) and ASMi (vertical stripes) for 16 h in a 96-well plate at 1% O_2_ with shaking. **C.** Comparison of WT and Δ*lasR* culture densities after growth in ASMi without and with KNO_3_ for 16 h at 1% O_2_ from panel A and B. Each data point represents an average of replicates from one day with lines connecting data from the same experiment. P-values were calculated using a paired one-way ANOVA with multiple comparisons (A and B), or a paired t-test (C).

### Dnr-regulated *norCB* were important for growth when KNO_3_ was present

Based on the lack of growth stimulation by nitrate in WT cultures (Figs. 3C and 4C), we speculated that nitrate consumption generated a toxic intermediate. To test this model, we first determined if *Δanr* or Δ*dnr* mutants showed evidence for nitrate utilization. Both the Δ*anr* and Δ*dnr* mutants consumed nitrate in LB after growth in 1% O_2_, but there was significantly less consumption by the Δ*anr*Δ*dnr* double mutant (Fig. 5A) indicating that either transcription factor could support nitrate consumption which is consistent with known redundancy in their regulation of genes involved in denitrification (18, 19, 22). The Δ*lasR*Δ*dnr* strain also consumed nitrate at both 21% and 1% O_2_ (Fig. S6), and the loss of *dnr* in the LasR- J215 strain did not affect nitrate consumption either (Fig. S4A). Previous work has shown that LasR- and Δ*lasR* mutants have higher Anr activity, higher expression of denitrification genes, and higher anaerobic denitrification rates (17). Interestingly, nitrate consumption of a Δ*lasR* mutant at 21% O_2_ was dependent on Anr as the supernatant of a Δ*lasR*Δ*anr* mutant had similar levels of nitrate as LB (P=0.079), and a Δ*lasR*Δ*anr*+*anr* strain consumed significant amounts of nitrate compared to LB (P<0.001) (Fig. S6). Although nitrate consumption was not dependent on Dnr, the Δ*dnr* mutant had lower relative growth in TB + 100, 200, 400 and 1600 µM KNO_3_ than the WT and the Δ*dnr*+*dnr* strains at the same concentrations when compared to their growth in TB alone (Fig. 5B). These data suggest that consumption of nitrate in the absence of Dnr activity may be toxic to *P. aeruginosa*.

**Figure 5.**
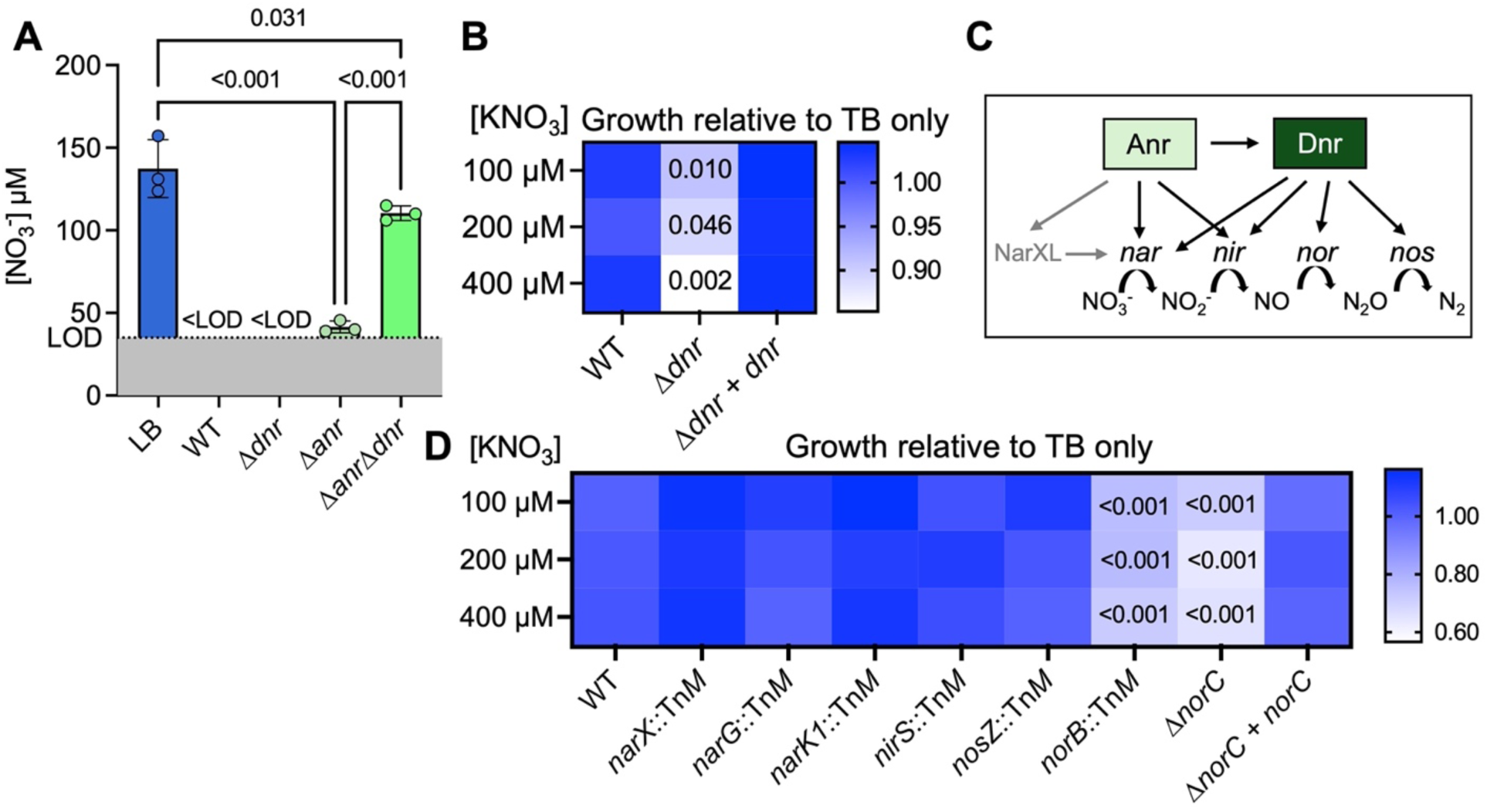
Nitrate consumption of *dnr* and *anr* mutants, and growth comparison of *P. aeruginosa* WT and transposon mutants defective in denitrification at different nitrate concentrations. **A.** The levels of NO_3_^-^ in LB before and after WT, Δ*dnr*, Δ*anr* and Δ*anr*Δ*dnr* growth for 16 h at 1% O_2_. NO_3_^-^ levels were normalized to OD_600_. Nitrate levels in LB are the same as in Figure 1B, and nitrate levels in WT supernatants at 1% O_2_ are the same as in Figure 1E. **B.** Growth of WT, a Δ*dnr* mutant, a Δ*dnr*+*dnr* strain in TB with indicated concentrations of KNO_3_ relative to growth in TB alone. Data in each cell represents the average of 9 experiments. For statistical analyses, relative growth of Δ*dnr* was compared to WT and Δ*dnr*+*dnr* at each concentration of KNO_3_ added. P-values were calculated using a paired one-way ANOVA with multiple comparisons. Only P-values of Δ*dnr* compared to WT are shown. **C.** A visual representation of the Anr- and Dnr-regulated dentification pathway. **D.** Relative growth of WT and confirmed PA14 Tn*M* mutants with insertions in specified genes, the Δ*norC* mutant, and the Δ*norC*+*norC* complemented strain. Colors represent growth at the specified [KNO_3_] divided by growth in TB without added KNO_3_. Cultures were grown in 96-well plates at 1% O_2_ for 16 sh on a shaker. Data in each cell represents the average of 3-5 experiments. For statistical analyses, relative growth of mutants was compared to WT at each concentration of KNO_3_ added. P-values were calculated using an ordinary one-way ANOVA with multiple comparisons. Only P-values <0.05 are shown.

To better understand the reason for the growth defect of Δ*dnr* strain concomitant with nitrate consumption, we analyzed the growth of a set of transposon mutants defective in denitrification outlined in Fig. 5C. The *narX*::Tn*M, narG*::Tn*M*, and *narK1*::Tn*M* mutants, which are defective in sensing, transport, and reduction of nitrate did not present with a decrease in relative growth upon the addition of nitrate in TB (Fig. 5D). After 16 h of growth at 1% O_2_, only the *nirS*::Tn*M* mutant had nitrite in its culture supernatants, and they were higher than those found in LB alone (P=<0.001) (Fig. S2). However, the WT and the *nirS* mutant grew similarly suggesting that nitrite accumulation was not inhibiting growth in the absence of Dnr activity (Fig. 5D). A *nosZ*::Tn*M*, defective in nitrous oxide reduction grew similarly to the no nitrate control across all nitrate concentrations tested (100-400 µM, Fig. 5D). In contrast, the *norB* transposon mutant (*norB*::Tn*M*) and the Δ*norC* in-frame deletion mutant showed reduced growth in TB with 100, 200 and 400 µM KNO_3_ while the Δ*norC*+*norC* strain grew similar to WT at each concentration of KNO_3_ (Fig. 5D). These data support a model in which NO reduction by nitric oxide reductase was important for growth in media with KNO_3_ and that regulation of *norCB* by Dnr and, in some settings Anr, contributes to NO resistance.

## Discussion

In this report, we showed that when oxygen is in the microoxic range, both Anr and Dnr transcription factors contributed to growth in media with micromolar concentrations of nitrate including in commonly used media such as LB (∼130 µM nitrate, Fig. 1B) and ASMi, a synthetic sputum medium with 340 µM KNO_3_. Our data suggest that nitrate consumption in oxic conditions did not enhance final yield (Figs. 3C and D and 4C). Rather, genetic analyses suggested that nitrate consumption generated inhibitory levels of the potentially toxic denitrification intermediate nitric oxide which necessitated the activities of Dnr and Dnr-regulated nitric oxide reductase (NorCB) (35–37) (Figs. 3D and 5B and D). *P. aeruginosa* Dnr is directly activated by nitric oxide upon its binding to a heme cofactor (24, 38–41). At atmospheric concentrations of O_2_ (21% O_2_), the WT strain did not consume nitrate (Fig. 1E) and the WT and Δ*dnr* mutant grew similarly (Fig. S3A). In contrast, strains defective in the quorum sensing regulator LasR (PA14 Δ*lasR* and clinical isolate J215 with a loss-of-function mutation in *lasR*) which have elevated activity of Anr even at atmospheric O_2_ (10), consumed nitrate at both 21% and 1% O_2_ (Fig. 2A and S4A), and required *dnr* for full growth at both oxygen tensions (Figs. S3B and S4B). Dnr was not required for fitness in 1% O_2_ in media without nitrate (Figs. 3A and B and 4A and B). These data emphasize the importance of Dnr even in low nitrate concentrations when denitrification is induced in response to oxygen limitation.

In denitrification, both nitrite (42, 43) and nitric oxide (44–46) intermediates have the potential for toxicity. However, the relative growth of mutants defective in different steps of denitrification showed that the *norB* and *norC* mutants, which accumulate nitric oxide, but not the *nirS* mutant, which accumulates nitrite (Fig. S2), had decreased fitness when compared to the WT strain. Previous studies showing that nitrite has growth inhibitory effects on *P. aeruginosa* (42, 43) were performed at nitrite concentrations of ∼15 mM which is much higher than the levels of nitrite in *nirS*::Tn*M* supernatants (∼30 µM) in LB-grown cultures (Fig. S2). Thus, we propose that the growth defect of a Δ*dnr* mutant in the presence of O_2_ is largely due to insufficient levels of nitric oxide reductase encoded by *norCB*). In infections, nitric oxide is toxic to pathogens including *P. aeruginosa* (47–49). In addition to the endogenous nitric oxide generated during denitrification, nitric oxide is produced as an antimicrobial agent by the innate immune system (50, 51). In fact, nitric oxide has been proposed as an antimicrobial therapy (48).

The requirement of *dnr* for full growth in ASMi at 1% O_2_ (Fig. 4A and B) supports other studies indicative of the potential for denitrification in infections such as those in the CF lung. Additionally, our work synthesizes studies that posit that denitrification and O_2_ respiration both occur in CF infections. For example, Dnr-regulated transcripts are high in *P. aeruginosa* RNA isolated from respiratory sputum (52), in cultured CF clinical isolates (53–55), and in cells grown in CF sputum (56, 57) and SCFM2 (29, 58). Dentrification supports anaerobic growth when nitrate is at the levels detected in CF sputum and present in SCMF2 (∼400 µM) (31, 36, 59), but in microoxic conditions, *P. aeruginosa* can also generate energy for growth using O_2_ for respiration (9). Thus, the observation that nitrate did not promote growth in microoxia with low levels of nitrate (Fig. 4C) is likely indicative of energy generation primarily through the respiration of O_2_. Considering the impacts and uses of NO, our studies highlight the importance of Dnr-regulated nitric oxide reductase activity in the CF infection environment.

The specific culture conditions may also influence the contribution of low concentrations of nitrate on *P. aeruginosa* growth. While our studies were performed in batch culture conditions, systems with a continuous input of nitrate may reveal a growth advantages from microoxic and, in LasR- strains, normoxic denitrification. The impact of denitrification in CF infections is likely variable as there is a range of nitrate concentrations measured in CF airway samples (30), patient status can affect O_2_ concentrations available to microbes (1), and strains differ in their capacity for denitrification when O_2_ is present (Fig. 2A). Thus, the relative contributions of denitrification and O_2_ respiration may change over space and time in a single person as well as between individuals.

Anr was necessary for full growth in microoxic conditions regardless of the presence of nitrate (e.g. Fig. 1C). Anr is important for *P. aeruginosa* virulence in murine lung infections (12) and in strains grown in CF sputum (57). Additionally, an Δ*anr* mutant had reduced growth in LB even at 21% O_2_ (Fig. 1A) while a Δ*dnr* mutant grew similarly to the WT (Fig. S3A). Anr regulates many genes that contribute to *P. aeruginosa* microoxic growth including the *ccoN2O2P2Q2* operon which encodes for a high-affinity *cbb*_3_-type terminal oxidase (60, 61), and *mhr* which encodes a hemerythrin that reversibly binds O_2_ with low micromolar affinities (10, 11). The expression of *hemN*, which encodes a protein necessary for O_2_-indepdendent heme biosynthesis (62) and genes involved in alternative energy generation such as *ldhA*, which encodes a lactate dehydrogenase and *arcDABC* operon which encodes the enzymes in the arginine deiminase pathway (63, 64) are also under Anr control. Anr also regulates *adhA*, which encodes an enzyme involved in the catabolism of exogenous ethanol, a fermentation product often made in O_2_-limited environments by species other than *P. aeruginosa*. (62, 65). Because NO can inactivate Anr (26, 46), the absence of Dnr activity may also limit Anr’s other roles. Together, these data suggest that both Anr and Dnr play important roles in the fitness at microoxic and normoxic conditions, and that the effects in normoxic conditions may be even stronger in LasR- strains and clinical isolates. These data may aid in the study of *P. aeruginosa* pathways relevant to disease, quorum sensing, and metabolism.

## Materials and methods

### Bacterial strains and growth conditions

All bacterial strains and plasmids used in this study are listed in Table S1. Bacteria were routinely grown in lysogeny broth (LB; 1% tryptone, 0.5% yeast extract, 0.5% NaCl) at 37 °C. Tryptone broth (TB; 1% tryptone, 0.5% NaCl) with or without the indicated concentrations of potassium nitrate (KNO_3_) and artificial sputum media for imaging (ASMi) and the same medium lacking the 340 µM KNO_3_ (ASMi-KNO_3_) were used for experiments where noted. The recipe for ASMi is described in (13).

### Construction of in-frame deletion and *att*::Tn7 site complementation mutants and plasmids

Primers used in plasmid construction are listed in Table S2. For the *norC* deletion construct, a gene block was ordered from Twist Biosciences and cloned into the pMQ30 allelic replacement vector. The *att*Tn7::*norC* complementation plasmid in which *norC* was expressed under its native promoter, was built using the T4 ligation protocol with T4 DNA ligase (New England Biolabs; M0202). Plasmids were confirmed by sequencing prior to introduction into *P. aeruginosa* by conjugation. Integration of the complementation construct was confirmed by PCR and restoration of anaerobic growth in LB with 100 mM nitrate to a Δ*norC* mutant.

### Growth assays

Cultures were inoculated from overnight cultures grown in 5 mL LB for 16 h that were normalized to OD_600_ = 1 in specified medium. For 96-well plate cultures, wells were inoculated to a starting OD_600_ = 0.05, then grown with aeration on a shake plate (Benchemark ORBi-SHAKER) at 225 rpm at 1% O_2_ or on a 96-well shaker (Thermo Labsystems Wellmix) set to 5 at 21% O_2_. Absorbance at 600 nm was read using a spectrophotometer (SpectraMax M2). For studies under microoxic conditions, cultures were grown inside a hypoxic cabinet with O_2_ and CO_2_ controllers (COY Laboratory Products, Grass Lake, MI), at 1% O_2_ and 5% CO_2_.

### Nitrate quantification and nitrate consumption assays

Nitrate quantification was performed using the API nitrate test kit according to the manufacturer’s protocol. In short, 2.5 mL of LB, TB, 0.5% yeast extract solution and 0.5% NaCl solution were added to glass test tubes. Next, 5 drops of solution I were added, and the solutions were mixed by vortexing briefly. Solution II was mixed by vigorously shaking for 30 s and then 5 drops were added to the solution I + media mixtures then the mixtures were vortexed for 1 min. After 5 min, 1 mL of the mixture was added to a 1 mL cuvette and the absorbance at 520 nm was read using a spectrophotometer (Thermo Genesys 6). Absorbance values were compared to a standard curve of potassium nitrate in H_2_O and used to calculate nitrate concentrations. Strains were grown at 37 °C in 5 mL tubes on either a roller drum at 21% oxygen or positioned diagonally in a test tube rack on a shake plate (Benchmark ORBi-SHAKER) at 225 rpm at 1% O_2_ for 16 h. Cells in the cultures were pelleted by centrifugation at 5000 rpm (Eppendorf 5804R, rotor A-4-44) in a 15 mL conical tube. The culture supernatant (2.5 mL) was transferred to a glass test tube for nitrate quantitation as described above.

### Nitrite quantification

Nitrite levels in culture supernatants were determined as previously described (66). In short, strains were grown at 37 °C in 5 mL tubes on either a roller drum at 21% oxygen or positioned diagonally in a test tube rack on a shake plate (Benchmark ORBi-SHAKER) at 225 rpm at 1% O_2_ for 16 h. Cells in the cultures were pelleted by centrifugation at 5000 rpm (Eppendorf 5804R, rotor A-4-44) in a 15 mL conical tube. The culture supernatant (1 mL) was transferred to a glass test tube and mixed with 1 mL of 0.02% *N*-(1-napthyl)ethylenediamine in 95% (v/v) ethanol and 1 mL of 1% sulfanilamide in 1.5 M hydrochloric acid and the absorbance at 550 nm was read using a spectrophotometer (Thermo Genesys 6). Absorbance values were compared to a standard curve of sodium nitrite in H_2_O and used to calculate nitrite concentrations.

### Validation of transposon mutants

To confirm the genomic location of the transposon insertion in mutants from the PA14 non-redundant collection (67), we performed arbitrary polymerase chain reaction (arbPCR) using primers listed in Table 2. Genomic DNA was isolated using MasterPure Yeast/bacteria Kit (Biosearch Technologies, #MPY80200) and diluted to 100 ng/µL. Primers were diluted to a concentration of 10 mM. The first round reaction mixture contained 2.5 µL 10X Standard Buffer (New England Biolabs, #B9014S), 1.5 µL of 50 mM magnesium chloride (MgCl_2_; New Englad Biolabs, #B0510A), 0.75 µL of PMFLGM.GB-3a, 1.5 µL of Arb1, 1.5 µL of Arb6, 5 µL of gDNA (100 ng/µL), 0.5 µL 10 mM dNTPs (PCR nucleotide mix, Roche chemicals; #63695222), 1.25 µL dimethyl sulfoxide (DMSO; Alfa Aesar, #36480), 0.3 µL Taq Polymerase (New England Biolabs, #M0273L), and 10.2 µL deionized water (dH_2_O) to bring up the volume to 25 µL. The following thermocycler (BioRad T100) protocol was used: after initial denaturation at 94 °C for 3 min, the reaction tubes were cycled 5 times at 94 °C for 30 sec, 30 °C for 30 sec, and 72 °C for 1 min with a 5 min extension at 72 °C. The second round reaction mix contained 2.5 µL 10X Standard Buffer (New England Biolabs, #B9014S), 1.5 µL of 50 mM magnesium chloride (MgCl_2_; New Englad Biolabs, #B0510A), 0.75 µL of PMFLGM.GB-2a, 1.5 µL of Arb2, 2µL of round 1 reaction mixture, 0.5 µL 10 mM dNTPs (PCR nucleotide mix, Roche chemicals; #63695222), 1.25 µL DMSO (Alfa Aesar, #36480), 0.3 µL Taq Polymerase (New England Biolabs, #M0273L), and 15.2 µL deionized water (dH_2_O) to bring up the volume to 25 µL. After initial denaturation at 94 °C for 3 min, the reaction tubes were cycled 30 times at 94 °C for 30 sec, 55 °C for 30 sec, and 72 °C for 1 min with a 5 min extension at 72 °C. The reactions were purified using QIAquick PCR Purification and Microcentrifuge Protocol (Qiagen, #28104). One µL of the purified products was sequenced with 1 µL of PMFLGM.GB-4a in a final volume of 20 µL of dH_2_O was analyzed by sequencing and the results were aligned to the strain PA14 genome.

### Statistics

Data analyses were performed using Graphpad Prism (version 10.3.0.

The specific statistical tests used are noted in each figure legend.

## Acknowledgements

The research reported in this publication was supported by R21 AI174132 grants from the National Institutes of Health (NIH) and the Cystic Fibrosis Foundation, GREEN19G0 and BOMBER24P0. Core support was provided by the NIDDK P30-DK117469 (Dartmouth Cystic Fibrosis Research Center or DartCF), BioMT (NIGMS P20GM113132) and the Dartmouth Molecular Biology Shared Resource (NCI 5P30CA023108). The transposon mutants used in this study were obtained from the PA14 non-redundant collection(67). We would like to thank Dr. Nicholas Jacobs (Dartmouth) and members of the Hogan Lab for helpful discussions.

---

**Figure S1.**
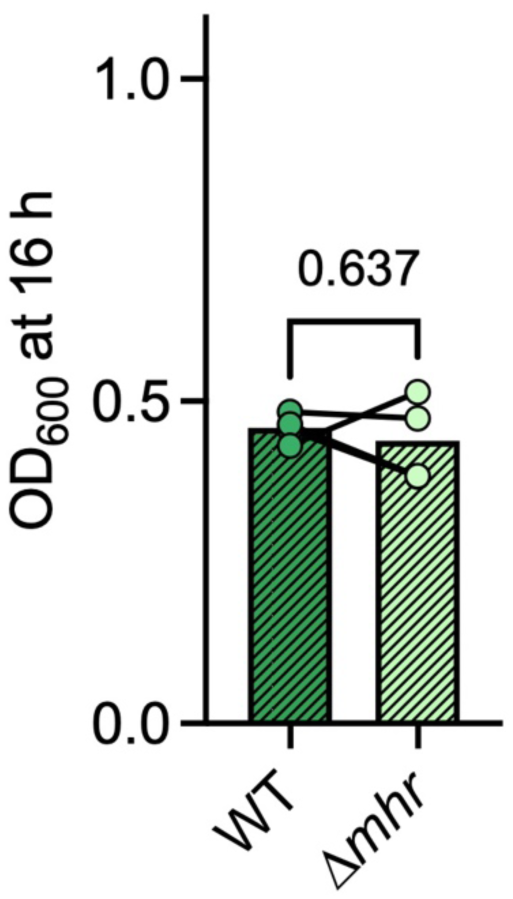
Anr-regulated *mhr* contribution to microoxic growth in LB. **A.** WT and Δ*mhr* growth at 1% O_2_ in LB in a 96-well plate for 16 h with shaking. Each data point represents an average of replicates from one day with lines connecting data from the same day. P-values were calculated using a paired t-test.

**Figure S2.**
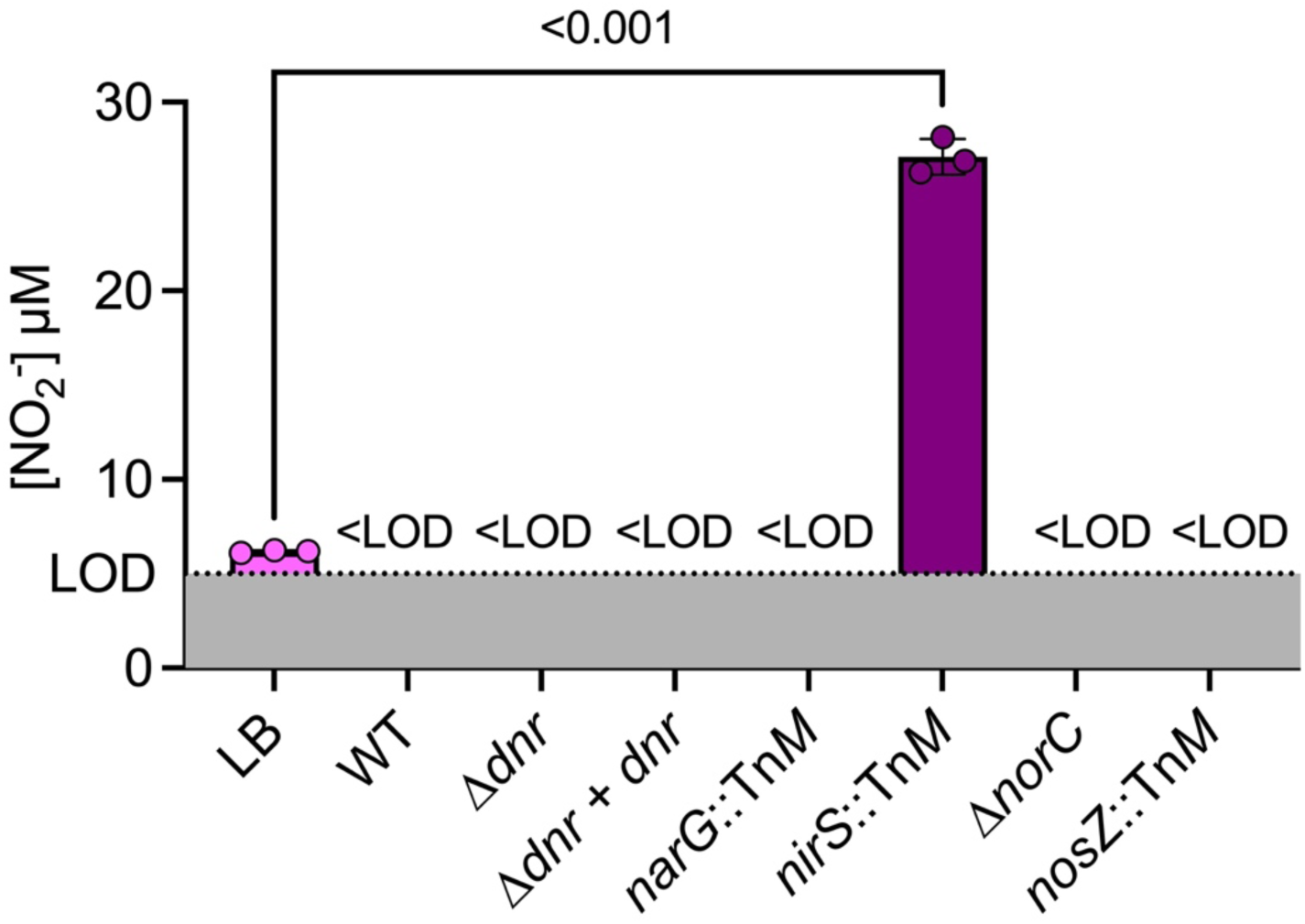
Levels of nitrite in LB and supernatants. Concentration of nitrite (NO_2_) in LB and in supernatants after 16 h of growth of indicated strains in 5 mL culture tubes. Concentrations were calculated using a standard curve of sodium nitrite NaNO_2_ in water. Levels below the limit of detection (LOD) are considered not detected (n.d.). Each data point represents a biological replicate, and the P-value was calculated using an unpaired t-test.

**Figure S3.**
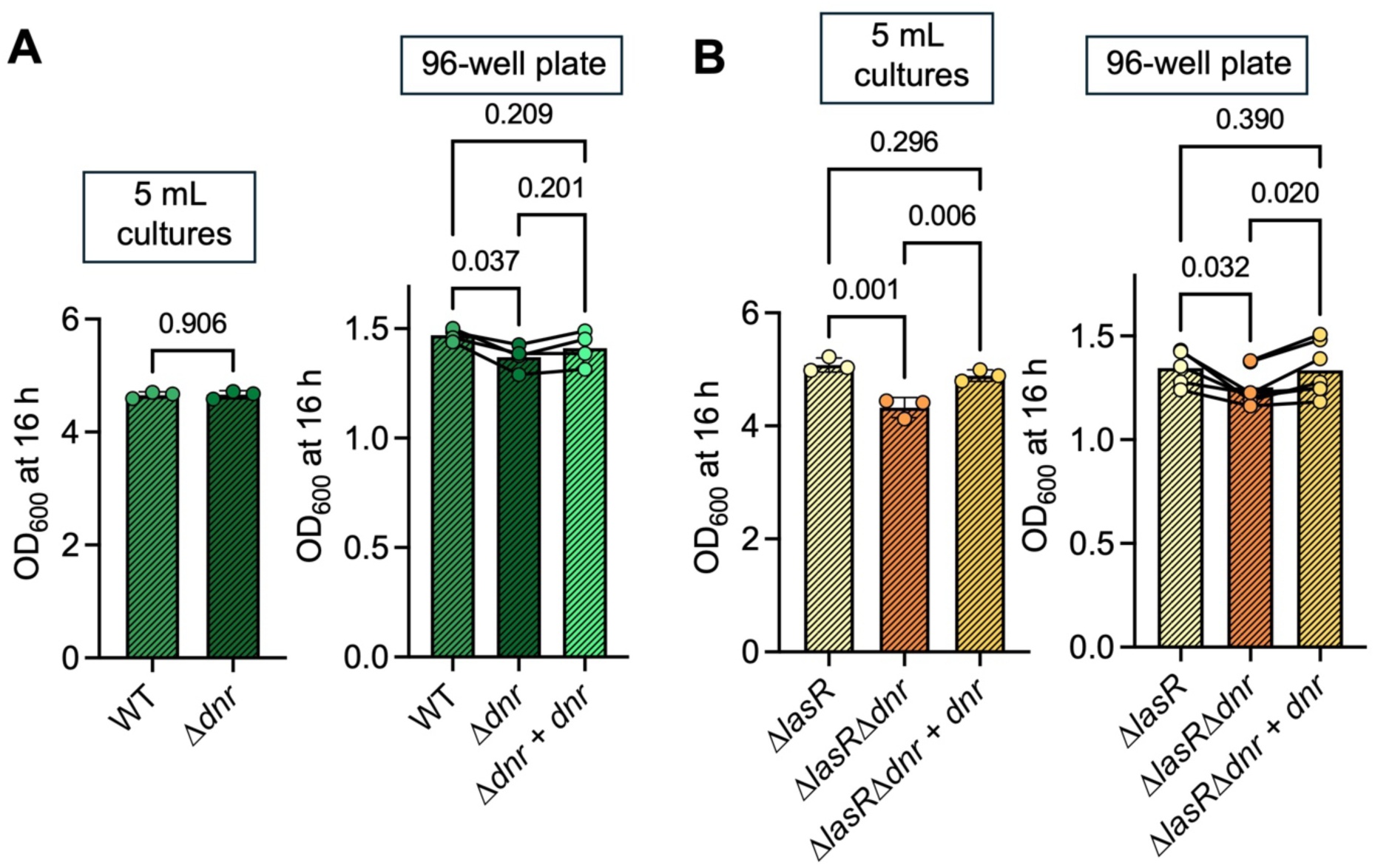
**Normoxic growth of *P. aeruginosa*. OD_600_ of WT, Δ*dnr*, and Δ*dnr*+*dnr*** (**A**) and Δ*lasR, ΔlasR*Δ*dnr* and Δ*lasR*Δ*dnr*+*dnr* (**B**) cultures grown in 5 mL LB on a roller drum, or 200 µL LB in a 96-well plate for 16 h on a shake plate at 21% O_2_. Data points from 5 mL cultures each represent a biological replicate, data points from 96-well plates represent an average of replicates from one day with lines connecting data from the same day. P-values were calculated using an unpaired t-test (A, 5 mL culutres), an ordinary one-way ANOVA with multiple comparisons (B, 5 mL cultures), or a paired one-way ANOVA with multiple comparisons (96-well plates).

**Figure S4.**
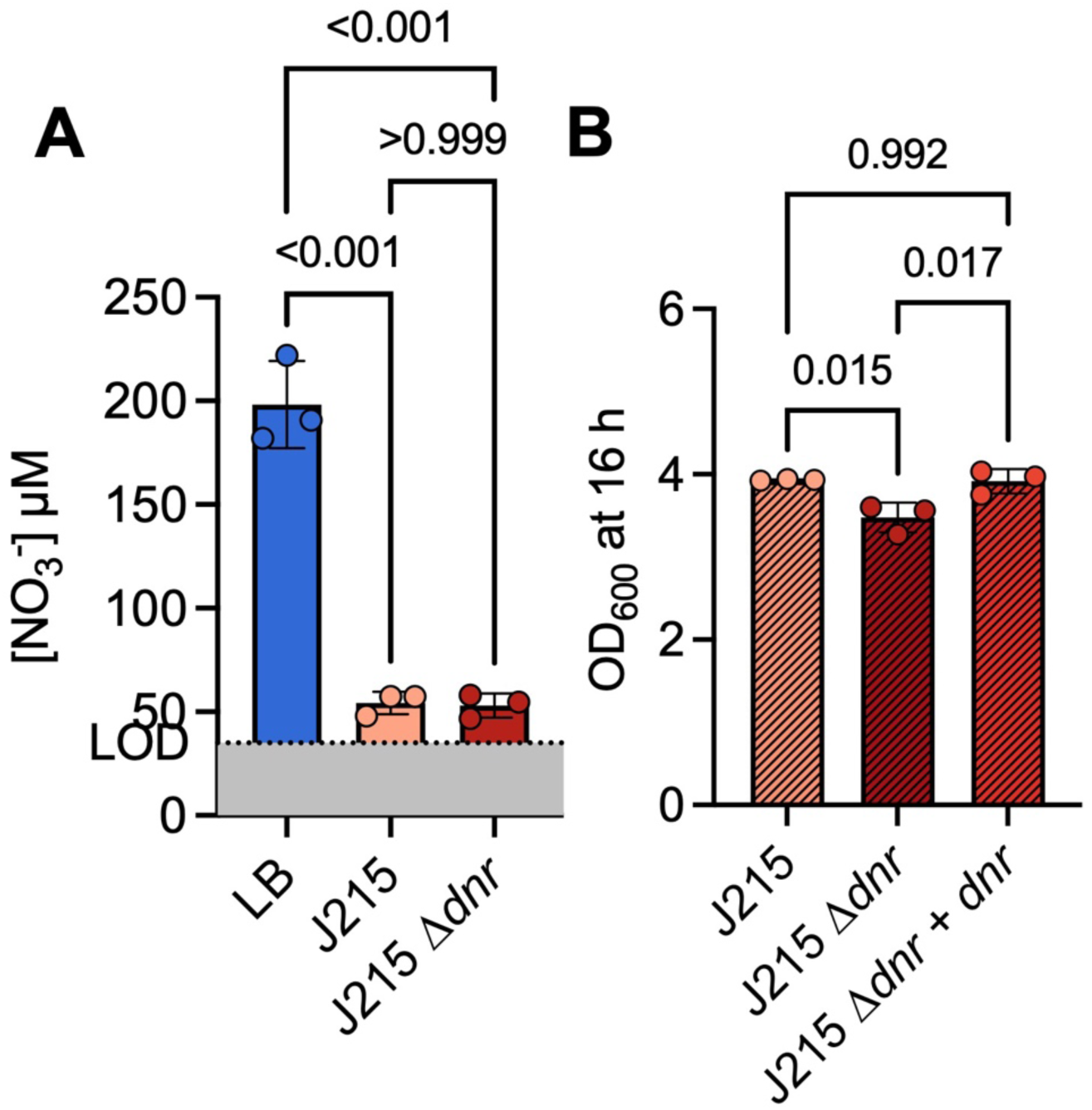
Nitrate consumption and growth of *P. aeruginosa* strain J215. **A.** The levels of nitrate in LB before and after 16 h of growth of the J215 strain and J215 Δ*dnr* mutant at 21% O_2_ for 16 h. NO_3_^-^ levels were calculated using a standard curve of KNO_3_ in water and normalized to OD_600_. Nitrate levels in LB are the same as in Figure 1B. **D.** Growth after 16 h of of J215, Δ*dnr* mutant and Δ*dnr*+*dnr* strain in 5 mL LB cultures. Data points each represent a biological replicate. P-values were calculated using an ordinary one-way ANOVA with multiple comparisons.

**Figure S5.**
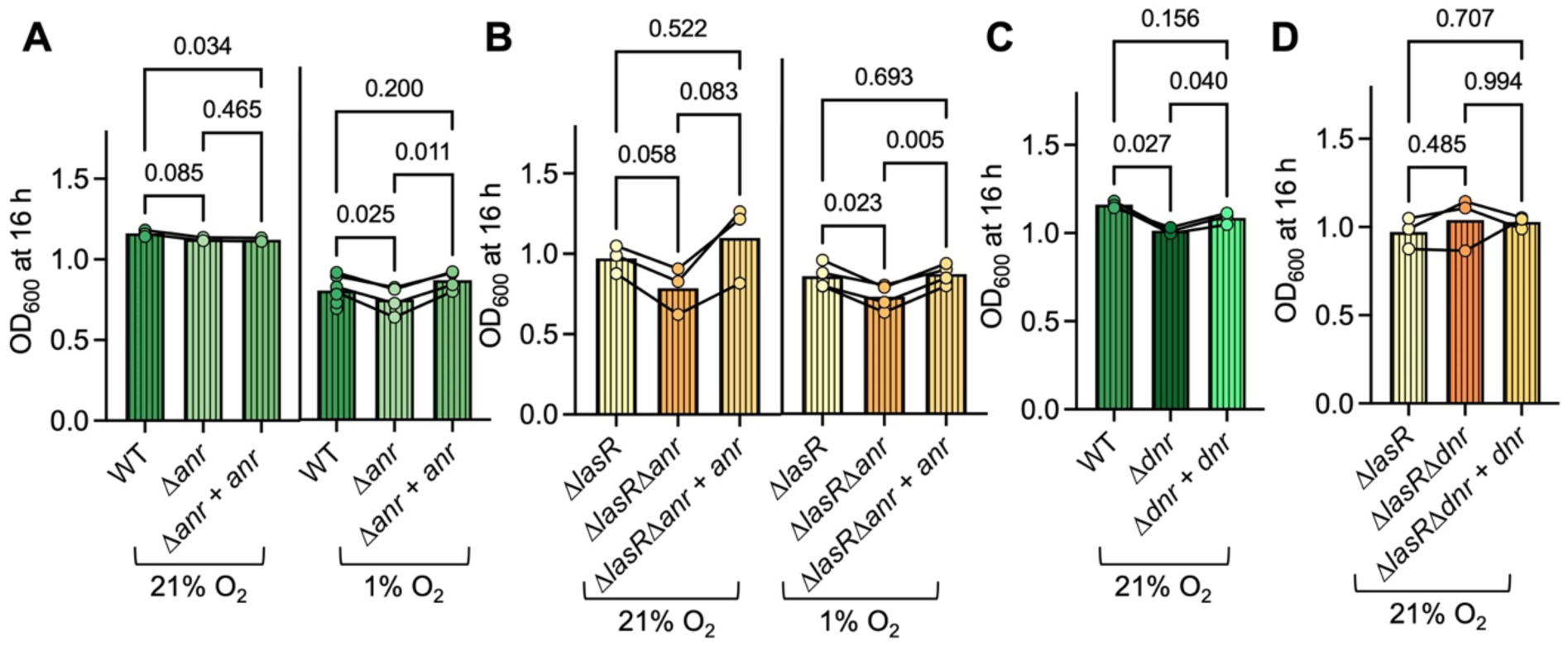
Anr- and Dnr-dependent growth of *P. aeruginosa* in ASMi. **A.** Culture densities for WT, Δ*anr* mutant, and the Δ*anr+anr* strain in ASMi at 21% and 1% O_2_ in a 96-well plate after 16 h. **B.** The culture densities of Δ*lasR*, Δ*lasR*Δ*anr*, and the Δ*lasR*Δ*anr+anr* strains grown in ASMi at 21% and 1% O_2_ in a 96-well plate for 16 h. **C.** Growth of WT, Δ*dnr* mutant, and Δ*dnr+dnr* strains in ASMi after 16 h in a 96-well plate at 21% O_2_. **D.** Culture densities of the Δ*lasR*, Δ*lasR*Δ*dnr* and the Δ*lasR*Δ*dnr+dnr* strain in ASMi in 96-well plates at 21% O_2_. All 96-well plates were grown with shaking. Each data point represents an average of replicates from one day with lines connecting data from the same day. All P-values were calculated using a paired one-way ANOVA with multiple comparisons.

**Figure S6.**
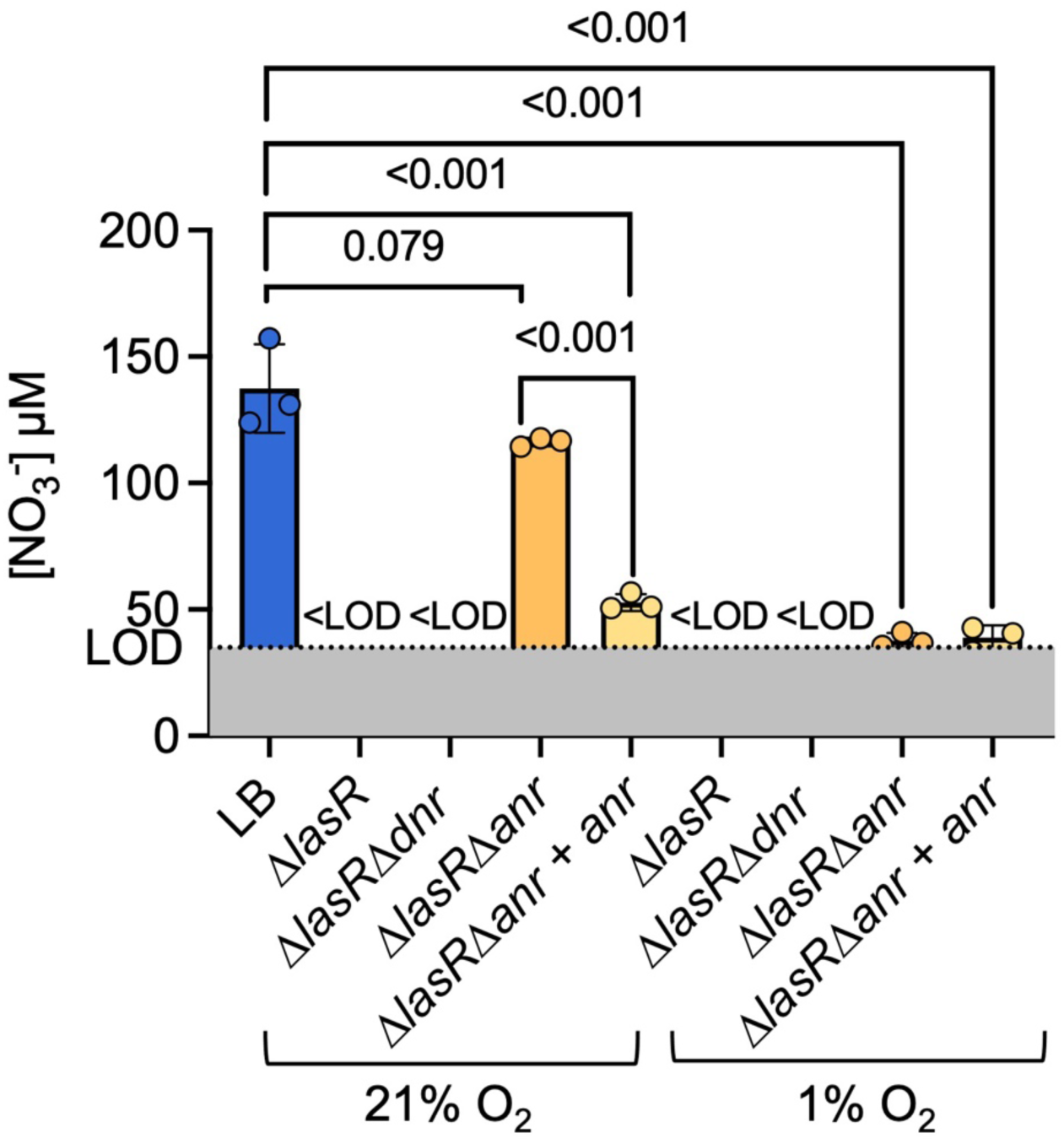
Nitrate consumption and growth of *P. aeruginosa* Δ*lasR* mutant. The levels of nitrate in LB before and after 16 h of growth of the Δ*lasR*, Δ*lasR*Δ*dnr*, Δ*lasR*Δ*anr* mutants and a Δ*lasR*Δ*anr*+*anr* strain at 21% and 1% O_2_ for 16 h. NO_3_^-^ levels were calculated using a standard curve of KNO_3_ in water and normalized to OD_600_. Levels of nitrate in LB are the same as in Figure 1B, levels of nitrate in Δ*lasR* supernatant at 21% and 1% O_2_ are the same as in Figure 2A. Each data point represents a biological replicate. P-values were calculated using an ordinary one-way ANOVA.

