## Supplemental Figures S1-S6 and Table S1 for "Role of *Pseudomonas aeruginosa* Dnr-regulated denitrification in oxic conditions"

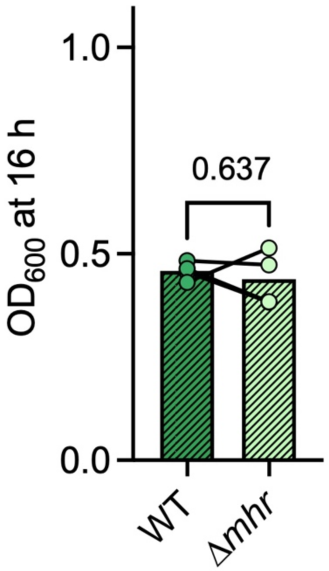


**Figure S1. Anr-regulated *mhr* contribution to microoxic growth in LB. A.** WT and Δ*mhr* growth at 1% $O_2$ in LB in a 96-well plate for 16 h with shaking. Each data point represents an average of replicates from one day with lines connecting data from the same day. P-values were calculated using a paired t-test.

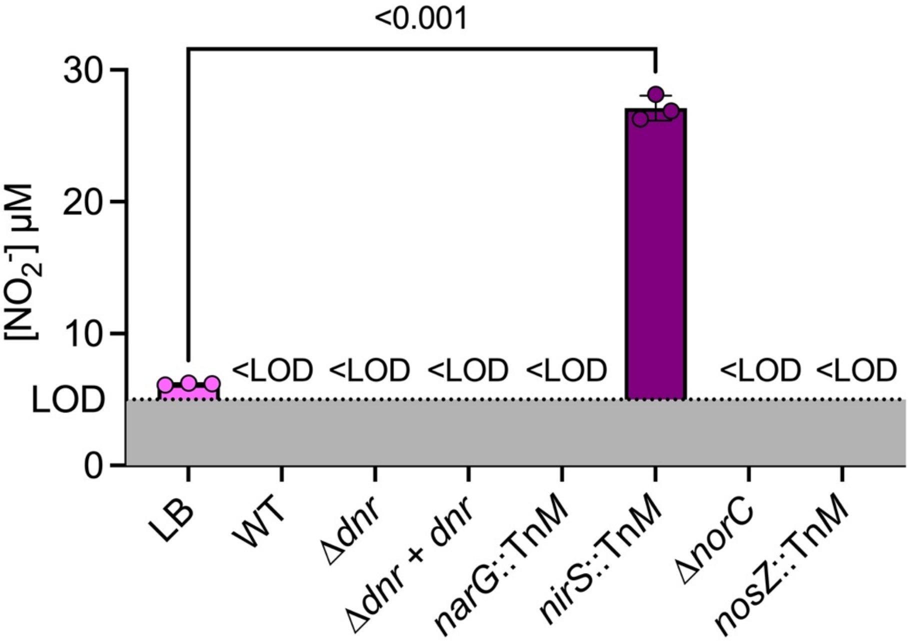


**Figure S2. Levels of nitrite in LB and supernatants.** Concentration of nitrite ($NO_2$) in LB and in supernatants after 16 h of growth of indicated strains in 5 mL culture tubes. Concentrations were calculated using a standard curve of sodium nitrite $NaNO_2$ in water. Levels below the limit of detection (LOD) are considered not detected (n.d.). Each data point represents a biological replicate, and the P-value was calculated using an unpaired t-test.

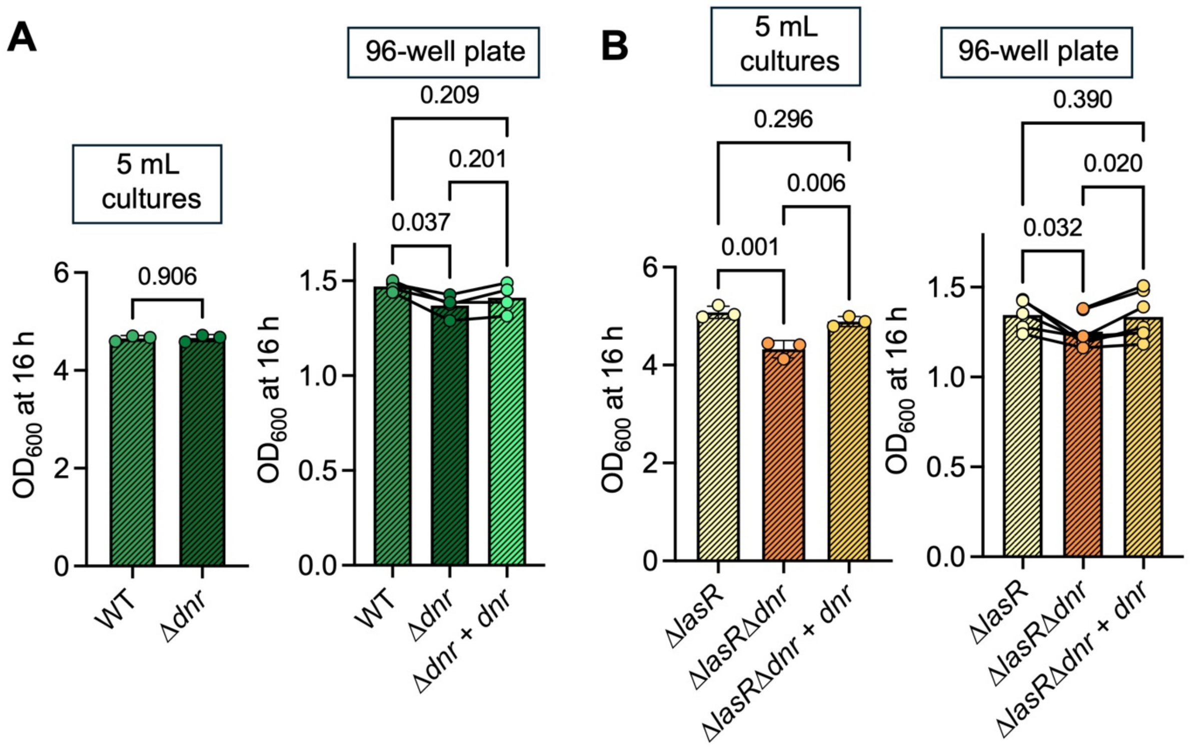


**Figure S3. Normoxic growth of *P. aeruginosa*.** $OD_{600}$ of WT, Δ*dnr*, and Δ*dnr*+*dnr* (**A**) and Δ*lasR,* Δ*lasR*Δ*dnr* and Δ*lasR*Δ*dnr*+*dnr* (**B**) cultures grown in 5 mL LB on a roller drum, or 200 µL LB in a 96-well plate for 16 h on a shake plate at 21% $O_2$. Data points from 5 mL cultures each represent a biological replicate, data points from 96-well plates represent an average of replicates from one day with lines connecting data from the same day. P-values were calculated using an unpaired t-test (A, 5 mL culutres), an ordinary one-way ANOVA with multiple comparisons (B, 5 mL cultures), or a paired one-way ANOVA with multiple comparisons (96-well plates).

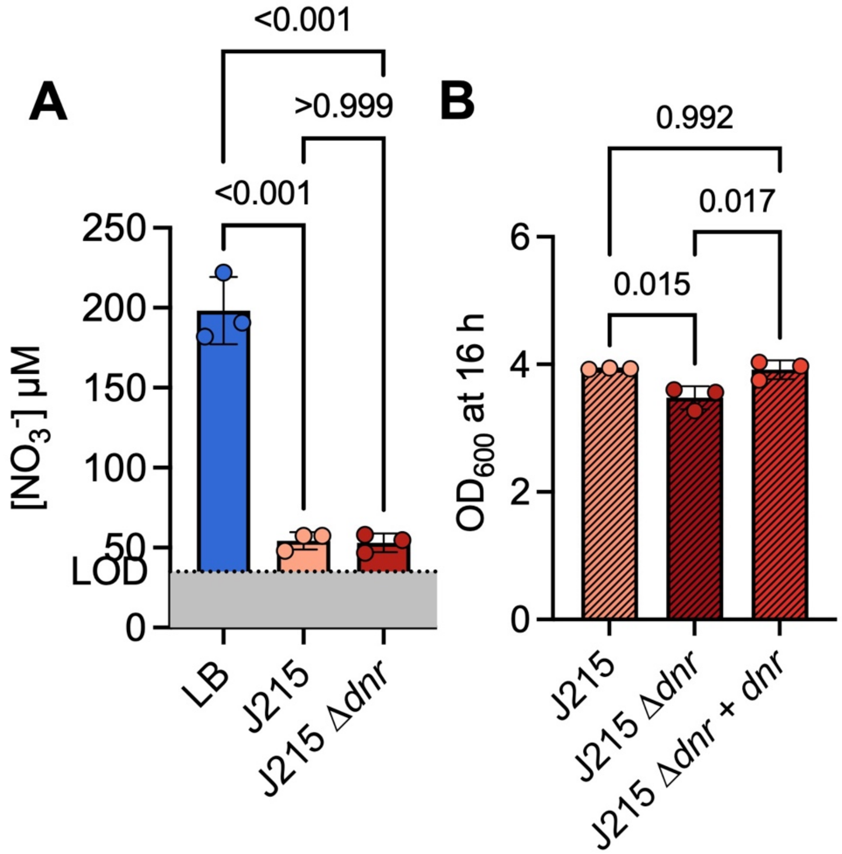


**Figure S4. Nitrate consumption and growth of *P. aeruginosa* strain J215. A.** The levels of nitrate in LB before and after 16 h of growth of the J215 strain and J215 Δ*dnr* mutant at 21% $O_2$ for 16 h. $NO_3^-$ levels were calculated using a standard curve of $KNO_3$ in water and normalized to $OD_{600}$. Nitrate levels in LB are the same as in Figure 1B. **D.** Growth after 16 h of of J215, Δ*dnr* mutant and Δ*dnr*+*dnr* strain in 5 mL LB cultures. Data points each represent a biological replicate. P-values were calculated using an ordinary one-way ANOVA with multiple comparisons.

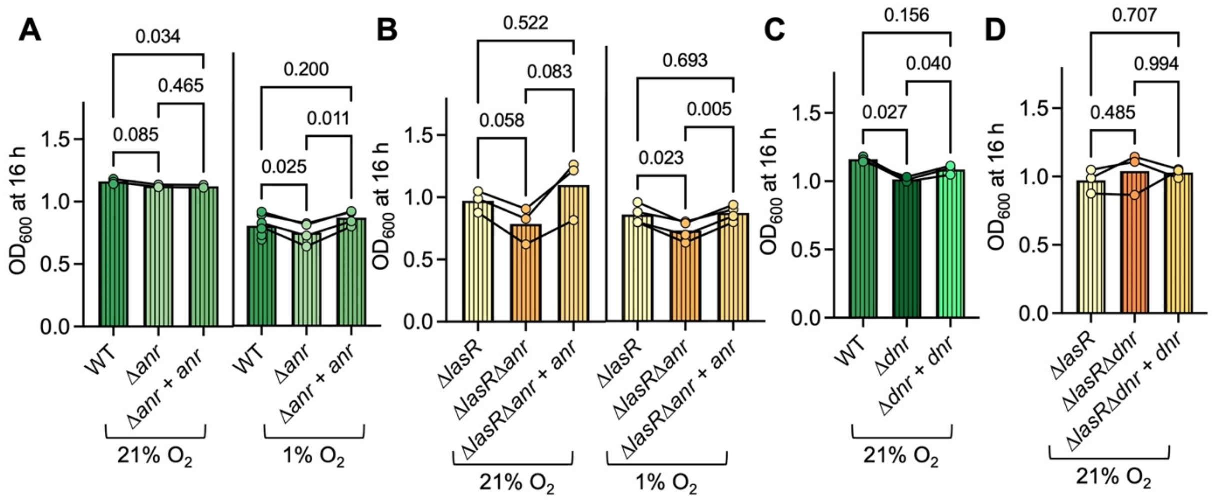


**Figure S5. Anr- and Dnr-dependent growth of *P. aeruginosa* in ASMi. A.** Culture densities for WT, Δ*anr* mutant, and the Δ*anr*+*anr* strain in ASMi at 21% and 1% $O_2$ in a 96-well plate after 16 h. **B.** The culture densities of Δ*lasR*, Δ*lasR*Δ*anr*, and the Δ*lasR*Δ*anr*+*anr* strains grown in ASMi at 21% and 1% $O_2$ in a 96-well plate for 16 h. **C.** Growth of WT, Δ*dnr* mutant, and Δ*dnr*+*dnr* strains in ASMi after 16 h in a 96-well plate at 21% $O_2$. **D.** Culture densities of the Δ*lasR*, Δ*lasR*Δ*dnr* and the Δ*lasR*Δ*dnr*+*dnr* strain in ASMi in 96-well plates at 21% $O_2$. All 96-well plates were grown with shaking. Each data point represents an average of replicates from one day with lines connecting data from the same day. All P-values were calculated using a paired one-way ANOVA with multiple comparisons.

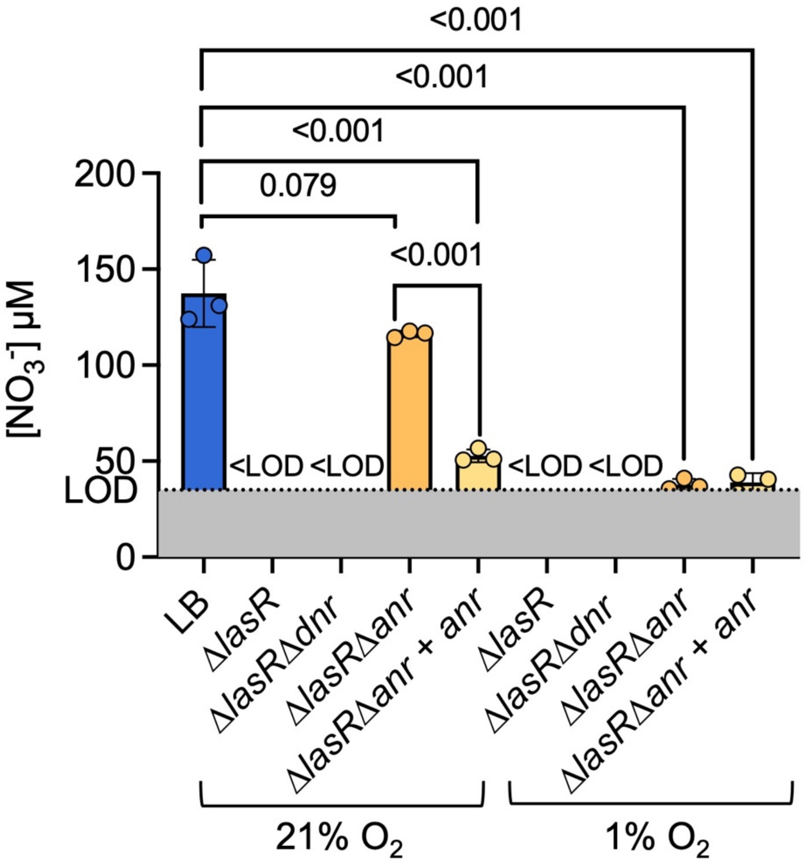


**Figure S6. Nitrate consumption and growth of *P. aeruginosa* Δ*lasR* mutant.** The levels of nitrate in LB before and after 16 h of growth of the Δ*lasR*, Δ*lasR*Δ*dnr*, Δ*lasR*Δ*anr* mutants and a Δ*lasR*Δ*anr*+*anr* strain at 21% and 1% $O_2$ for 16 h. $NO_3^-$ levels were calculated using a standard curve of $KNO_3$ in water and normalized to $OD_{600}$. Levels of nitrate in LB are the same as in Figure 1B, levels of nitrate in Δ*lasR* supernatant at 21% and 1% $O_2$ are the same as in Figure 2A. Each data point represents a biological replicate. P-values were calculated using an ordinary one-way ANOVA.

**Table S1. Strains and plasmids used in this study**

| Strain or plasmid | Strain ID | Description | Source |
|---|---|---|---|
| ***P. aeruginosa*** | | | |
| PA14 WT | DH122 | Wild-type *P. aeruginosa* | (1) |
| Δ*anr* | DH2855 | DH122 with in frame deletion of *anr* (PA14_44490) | (2) |
| Δ*anr*+*anr* | DH3478 | DH2855 with complementation of *anr* (PA14_44490) at the native locus | (2) |
| Δ*dnr* | DH1978 | In-frame deletion of *dnr* | |
| Δ*dnr att*Tn7::*dnr* | DH4598 | DH1978 with complementation of *dnr* at the Tn*7* attachment (*att*Tn7) site, GmR | This study |
| Δ*lasR* | DH164 | DH122 with in frame deletion of *lasR* (PA14_45960) | (3) |
| Δ*lasR*Δ*anr* | DH2401 | In frame deletions of *lasR* (PA14_45960) and *anr* (PA14_44490) | (4) |
| Δ*lasR*Δ*anr*+*anr* | DH3479 | DH2401 with complementation of *anr* at the native locus | (5) |
| Δ*lasR*Δ*dnr* | DH3813 | In-frame deletion of *lasR* (PA14_45960) in DH1978 | This study |
| Δ*lasR*Δ*dnr* *att*Tn7::*dnr* | DH4599 | DH3813 with complementation of *dnr* at the *att*Tn7site, GmR | This study |
| J215 | DH2403 | Clinical isolate J215 (with a non-functional LasR) | (4) |
| J215Δ*dnr* | DH2409 | DH2403 with in frame deletion of *dnr* (PA14_45960) | This study |
| J215Δ*dnr* *att*Tn7::*dnr* | DH4602 | DH2409 with complementation of *dnr* at the *att*::Tn*7* site, GmR | This study |
| *narG*::Tn*M* | DH1754 | Transposon insertion in *narG* (gene), GmR | (6) |
| *narK1*::Tn*M* | DH1755 | Transposon insertion in *narK1* (gene), GmR | (6) |
| *narK*2::Tn*M* | DH1756 | Transposon insertion in *narK2* (gene), GmR | (6) |
| *narX*::Tn*M* | DH1757 | Transposon insertion in *narX* (gene), GmR | (6) |
| *nirS*::Tn*M* | DH1758 | Transposon insertion in *nirS* (gene), GmR | (6) |
| *norB*::Tn*M* | DH4601 | Transposon insertion in *norB* (gene), GmR | (6) |
| Δ*norC* | DH4664 | DH122 with in frame deletion of *norC* (PA14_06810) | This study |
| *nosZ*::Tn*M* | DH1760 | Transposon insertion in *nosZ* (gene), GmR | (6) |
| Δ*norC* *att*Tn7::*norC* | DH4750 | DH4664 with in frame complementation of *norC* at the *att*::Tn7 site, GmR | This study |
| ***E. coli*** | | | |
| SM10 + mini Tn*7*-*dnr* | DH4600 | *E. coli* strain SM10 containing the plasmid to complement *dnr* for conjugation into *P. aeruginosa*, GmR | (7) |

| | | | |
|---|---|---|---|
| S17 + mini Tn7-*norC* | DH4749 | *E. coli* strain S17 containing the plasmid to complement *norC* for conjugation into *P. aeruginosa*, GmR | This study |
| S17 + pMQ30_Δ*norC* | DH4743 | *E. coli* strain S17 containing the plasmid for in-frame deletion of *norC* (PA14_06810) | This study |
| **Plasmids** | | | |
| puc18T-mini-Tn*7*T-Gm | | *att*Tn*7* site insertion vector, GmR | (8) |
| puc18T-mini-Tn*7*-*dnr* | | plasmid to complement *dnr*, with *dnr* under the control of its native promoter, at the *P. aeruginosa att*Tn*7* site, GmR | (7) |
| puc18-mini-Tn7-*norC* | | Plasmid to complement *norC*, with *norC* under the control of its native promoter, at the *att*Tn*7* site in *P. aeruginosa*, GmR | This study |
| pMQ30 | | Allelic replacement vector, GmR | (9) |
| pMQ30_Δ*norC* | | Plasmid for in-frame deletion of *norC*, GmR | This study |

**Table S2. Primers used in this study**

| Primer | Primer No. | Description |
|---|---|---|
| Arb1 | DH179 | Arbitrary primer 1 (5'-GGCCACGCGTCGACTAG TACGGNNNNNNNNNNGATAT-3') |
| Arb2 | DH180 | Arbitrary primer 2 (5'-GGCCACGCGTCGACTAG TACNNNNNNNNNNACGCC-3') |
| Arb6 | DH181 | Arbitrary primer 6 (5'-GGCCACGCGTCGAC TAGTACGGNNNNNNNNNNACGCC-3') |
| PMFLGM.GB-3a | DH115 | Tn*M*-specific primer Round 1 (5'-TACAGTTTA CGAACCGAACAGGC-3') |
| PMFLGM.GB-2a | DH114 | Tn*M*-specific primer Round 2 (5'-TGTCAACTG GGTTCGTGCCTTCATCC-3') |
| PMFLGM.GB-4a | DH116 | Tn*M*-specific sequencing primer (5'-GACCGAGATAGGGTTGAGTG-3') |
| norC_down_HindIII | SB1 | Amplification of *norC* for cloning into puc18T-mini-Tn7 (5'-GAGACAAGCTTCAACCCTCCTTGTTCGGCG GC-3') |
| norC_up_BamHI | SB2 | Amplification of *norC* for cloning into puc18T-mini-Tn7 (5'-GAGACGGATCCGTGGCCAAGGGCTGGCTGA TC-3') |
